# Understanding genetic changes underlying the molybdate resistance and the glutathione production in *Saccharomyces cerevisiae* wine strains using an evolution-based strategy

**DOI:** 10.1101/092007

**Authors:** Francesco Mezzetti, Justin C. Fay, Paolo Giudici, Luciana De Vero

**Author notes:** **Author Contributions:** Conceived and designed the experiments: FM JCF LDV. Performed the experiments: FM. Analyzed the data: FM JCF LDV. Contributed reagents/materials/analysis tools: JCF PG. supervision: JCF PG LDV. Wrote the paper: FM LDV. Revised the original draft: LDV JCF PG.

## Abstract

In this work we have investigated the genetic changes underlying the high glutathione (GSH) production showed by the evolved *Saccharomyces cerevisiae* strain UMCC 2581, selected in a molybdate-enriched environment after sexual recombination of the parental wine strain UMCC 855. To reach our goal, we first generated strains with the desired phenotype, and then we mapped changes underlying adaptation to molybdate by using a whole-genome sequencing. Moreover, we carried out the RNA-seq that allowed an accurate measurement of gene expression and an effective comparison between the transcriptional profiles of parental and evolved strains, in order to investigate the relationship between genotype and high GSH production phenotype.

Among all genes evaluated only two genes, *MED2* and *RIM15* both related to oxidative stress response, presented new mutations in the UMCC 2581 strain sequence and were potentially related to the evolved phenotype.

Regarding the expression of high GSH production phenotype, it included over-expression of amino acids permeases and precursor biosynthetic enzymes rather than the two GSH metabolic enzymes, whereas GSH production and metabolism, transporter activity, vacuolar detoxification and oxidative stress response enzymes were probably added resulting in the molybdate resistance phenotype. This work provides an example of a combination of an evolution-based strategy to successful obtain yeast strain with desired phenotype and inverse engineering approach to genetic characterize the evolved strain. The obtained genetic information could be useful for further optimization of the evolved strains and for providing an even more rapid approach to identify new strains, with a high GSH production, through a marked-assisted selection strategy.

## Introduction

Evolutionary engineering or adaptive laboratory evolution is a powerful approach, widely used for improving industrially significant *Saccharomyces cerevisiae* strains. In the oenological field, it has been successful in engineering several phenotypes by generating wine yeast strains with improved properties, such as enhanced substrate utilization, tolerance to fermentation conditions and resistance to toxic compounds [1–5].

The adaptive evolution technique, which simply mimics nature by random mutation of the microorganisms’ own genes followed by selection under suitable conditions to favor the desired phenotype, has the main advantage of not requiring a priori knowledge of the genes involved in the expression of desired phenotypes [6–8].

However, direct evolutionary strategies can suffer from being time-consuming since extensive cultivation periods and multiple rounds of screening are often required [9]. Indeed, when the selection is non-targeted and non-specific for the type of mutation or not based on growth-linked traits a large number of strains need to be screened for isolating an improved mutant from a mixed population [10]. Moreover, the screening could be hindered by the difficulty in achieving evolved variants expressing selectable phenotypes [11].

To overcome this limit, novel evolutionary strategies have been designed for generating phenotypic diversity in a strains' population, and to develop cultivation strategies that effectively select cells with desirable phenotypes. Among them, the use of anti-metabolites or metabolite analogs, as selective pressures, have already been applied in evolutionary engineering to either improve productivity or improve substrate utilization [8].

Accordingly, we have designed an evolution-based strategy useful for the selection of *S. cerevisiae* wine strains that produce low levels of SO_2_ and H_2_S and high levels of GSH by using toxic sulfate analogues such as chromate Cr(VI) or molybdate Mo(VI) [11,12].

These heavy metals enter a yeast cell through high-affinity sulfate permeases and are involved in the sulfate assimilation as well as GSH biosynthetic pathway. Therefore, the resulting resistant strains are potentially able to show the desired phenotype being impaired in this metabolic pathway.

To outline our strategy, first sexual recombination and mating were used to increase strain randomization and second, a specific selective pressure was defined for generating the preferred recombinants by targeting the biosynthetic pathway of the metabolite of interest. Finally, the evolved strains potentially capable of expressing the desired phenotype were screening to select the best candidate.

Recently, by applying this evolution-based strategy, we have obtained the strain UMCC 2581 resistant to Mo(VI) 5 mM and with enhanced GSH content, at the end of the fermentation in a microvinification assay, with an increase of 100% compared to the parental strain UMCC 855 [12].

This achievement has confirmed the effectiveness of both meiotic recombination in generating clones with different and frequently better properties than their parental strain [13,14], and application of targeted selective pressure for obtaining candidate strains for the phenotype of interest. Above all, the exploitation of the Mo(VI) resistance has proved to be useful for the selection of the desired evolved strains, probably by activating the yeast common metal response that involves sulfur assimilation and GSH biosynthesis [12].

Nevertheless, the exact mechanism of resistance inside the cells had not been explained, therefore, a genetic characterization of the evolved and parental strains, in particular regarding the genes associated with GSH metabolism, is required for understanding their different behavior on a molecular level.

In yeast cells as well as other microorganisms, stress responses are affected by rapid adjustment of gene expression patterns [15].

Generally, the genetic dynamics found in wild wine yeast strains as well as in evolved strains vary in a multi-factor continuous way, and result from a wide assortment of evolutionary forces, among which mutation, selection, recombination and drift [16–18]. Thus, the application of the current high-throughput DNA and RNA sequencing (RNA-seq) technologies is extremely useful to achieve a rapid identification of genetic variations facilitating an efficient genetic mapping. In particular, RNA-seq has proved to be a better method for the study of industrial wine strains compared to microarrays [19]. Certainly, it is more sensitive in detecting genes with very low expression and more accurate in the quantification of highly expressed genes, due to its wider dynamic range [20,21].

In the present work, we intended to integrate our evolution-based strategy with an inverse engineering approach. We first generated strains with the desired phenotype by using a random method and then, through analysis of the genetic and transcriptomic of the evolved strains, mapped changes underlying adaptation to molybdate and the relationship between genotype and high GSH production phenotype. To reach this goal, we combined whole-genome sequencing, which reveals the repertoire of point mutations and copy number polymorphisms, with the gene expression analysis carried out by RNA-seq that allows an accurate measurement of gene expression and an effective comparison between the transcriptional profiles of parental and evolved strains.

In particular, we analyzed the molybdate resistance phenotype as a specific quantitative trait, in order to understand the relations with and the reasons of the higher GSH production showed by the evolved strain UMCC 2581 in comparison to UMCC 855 parental strain.

## Materials and methods

### Yeast strains and growth conditions

The *Saccharomyces cerevisiae* parental strain UMCC 855 and the evolved strain UMCC 2581 used in our experiments are described in Table 1.

**Table 1.** Parental and evolved *Saccharomyces cerevisiae* strains

| UMCC<sup>a</sup> code | Other name | Description | Reference |
| --- | --- | --- | --- |
| UMCC 855 | 21T2 | Laboratory yeast strain selected for its oenological aptitude. | [12,22,23] |
| UMCC 2581 | Mo21T2-5 | Evolved yeast strain from UMCC 855, high GSH producer. | [12] |

<sup>a</sup>Unimore Microbial Culture Collection (UMCC), University of Modena and Reggio Emilia- Reggio Emilia (Italy)

Strain UMCC 855 was used to generate the monosporic clones (MCs) throughout this study. Yeast cells were grown at 28 °C on YPD complete medium (1% yeast extract, 2% peptone and 2% glucose) with the addition of 2% agar when necessary. The strains are deposit in the Unimore Microbial Culture Collection (University of Modena and Reggio Emilia- Reggio Emilia - Italy) and stored for long-term preservation at -80 °C in cryovials containing YPD medium supplemented with 25% glycerol (v/v) as cryopreservative.

### Generation and screening of Molybdate-resistant monosporic clones

An overnight culture of UMCC 855 grown at 28 °C on YPDA, was resuspended in 3 mL of 1% Potassium Acetate (Fisher Scientific) and incubated at 28 °C overnight with shaking at 300 rpm to induce sporulation. The culture containing asci were diluted 2-fold with a solution of 10 mg mL^−1^ Zymolyase (20T, Fisher Scientific) and spotted on YPDA plates successively incubated for 1 h at 28 °C. After tetrad dissection, performed using a micromanipulator (Singer Instruments MSM System 200), the YPDA plates were incubated again at the same previous conditions until the growth of segregantes was observed. The MCs obtained were screened on YNB minimal medium supplemented with Mo(VI) at the concentrations of 0 (control plates), 1.0, 2.5, and 5.0 mM to evaluate their resistance phenotype according to Mezzetti et al. [12] (S1 Fig). Colony growth was observed after 4 days of incubation at 28 °C.

### Genomic DNA extraction, sequencing and data handling

Genomic DNA (gDNA) was extracted using the ZR fungal/bacterial DNA miniprep kit (Zymo Research) following the manufacturer’s instructions. The gDNA of the MCs belonging to the two clusters selected were pooled in equimolar amount and the whole-cluster gDNA was sequenced along with gDNA of the parental and evolved strains.

Genome sequencing was performed with the Ion Torrent ProtonTM platform (Life Technologies) by the Genomics Core Facility at Saint Louis University School of Medicine. Sequence reads were mapped to the *S. cerevisiae* reference genome, S288c [24], using BWA (Burrows-Wheeler Aligner) [25]. Duplicate sequences were removed using PicardTools 1.114 (http://broadinstitute.github.io/picard) and the Genome Analysis Toolkit (GATK v3.4-46) [26] was used for the variant discovery (single nucleotide polymorphisms, SNPs; insertions and deletions, InDels). The information was finally saved in “.vcf” format (variant call format).

In order to map the Quantitative Trait Loci (QTL) involved in the evolved phenotype, we selected the heterozygous sites of the parental strain UMCC 855 using a self-written perl script on the list in “vcf” format. The Alleles Frequencies (AF) were calculated and only sites with an AF ranging from 0.25 - 0.75 and coverage of greater than 20 reads were used in the subsequent comparison. In the comparison of Clusters Resistant-Parental/Resistant-Evolved a p-value was assigned to each site using Fisher’s Exact Test. A −log_10_(p-value) greater than 20 log unit was chosen as a cutoff to call QTL peaks and the width of each peak was determined by dropping 5 log units.

The SNPs and InDels specific to the parental strain UMCC 855 and evolved strain UMCC 2581 were identified from the vcf file. The discovered variants were analyzed using the snpEff software [27] (http://snpeff.sourceforge.net) to classify them according to their effect on protein-coding genes. Reads alignments and subsequent comparison of the “vfc” files underwent a strict quality and coverage verification that led to the exclusion of SNPs and InDels with coverage lower than 20x and frequency lower than 25% of the reads.

To discover regions of chromosomal copy-number variations (CNVs), the average sequencing coverage over a 1000 bp window size were calculated using IGVtools [28].

### RNA-sequencing and differential gene expression analysis

To capture gene expression changes between the parental and evolved strains we performed RNA-seq experiments. The two strains were inoculated in Erlenmeyer flasks filled with 100 mL of chemically defined synthetic must (SM) prepared according to Giudici & Kunkee [29]. Cell growth was monitored by measuring the optical density at 600 nm hourly until reaching the end of exponential phase. At each time point, the cells were harvested and centrifuged, then supernatant was removed and the pellet was immediately frozen in liquid nitrogen and stored at -80 °C until sample analysis. For each sample, the time point corresponding to three quarters through the exponential phase was chosen and the total RNA was extracted with hot acidic phenol [30]. mRNA purification, RNA-Seq library preparation and sequencing were performed by the Genomics Core Facility at Saint Louis University School of Medicine using Ambion Dynabeads mRNA Direct Micro Kit (Life Technologies), Ion Total RNA-seq kit v2 (Life Technologies) and the Ion Torrent ProtonTM platform (Life Technologies) respectively. Bowtie 2 [31] was used to align reads for each sample to the reference genome S288c, Picard 1.114 to eliminate duplicate reads and DESeq [32] was used to identified genes differentially expressed between the parental and evolved strains. Only genes with a false discovery rate (FDR) lower than 0.05 were considered differentially expressed. The gene ontology analysis was performed by using the on-line tool GO Term Finder (http://www.yeastgenome.org/cgi-bin/GO/goTermFinder.pl) within the SGD database. The statistical analysis of the GO term “molecular function” was performed using the BiNGO plug-in [33] in Cytoscape (http://www.cytoscape.org).

### Accession number

All DNA sequencing and RNA-seq data are available from the NCBI Sequence Read Archive (SRA, https://www.ncbi.nlm.nih.gov/sra/) with accession number SRP094104.

## Results

### QTL mapping

To study the genetic basis resistance to Mo(VI) in the evolved strain, we used a QTL mapping approach (Fig 1), developed in a three steps process similar to that proposed by Parts et al. [34]. First, we generated a pool of 69 MCs (which have been coded from UMCC 2665 to UMCC 2733) starting from parental strain UMCC 855. Then, we screened them on the YNB selective media, with the addition of different molybdate Mo(VI) concentrations according to our previous work [12]. The segregation analysis revealed a large amount of phenotype variation in the progeny, indicating the parental strain carried heterozygous mutations affecting resistance to Mo(VI).

**Fig 1.**
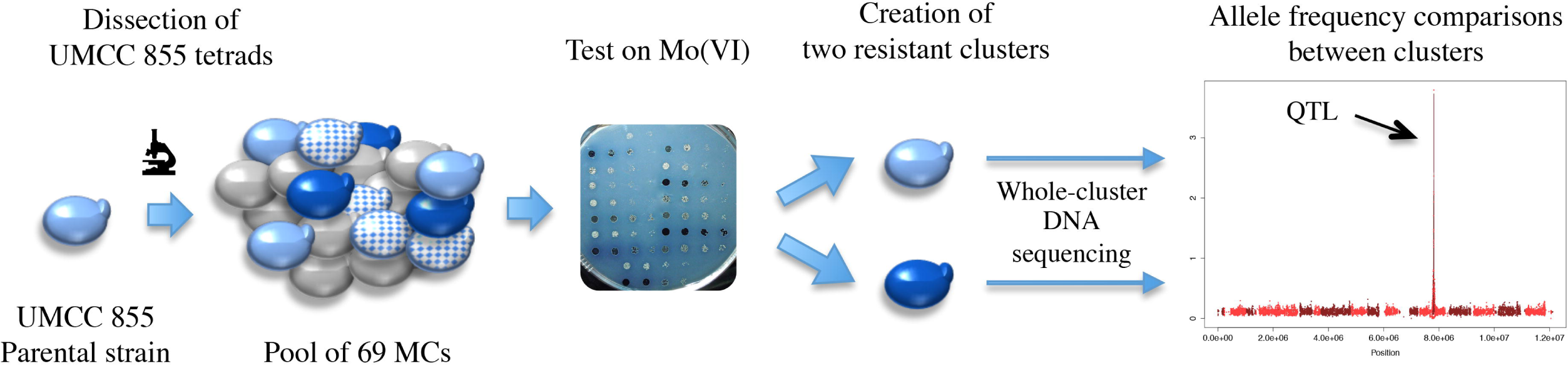
QTL mapping strategy. Creation of pool of spores by dissection of UMCC 855, test on selective media, formation of cluster according to the phenotype and detection of fixed alleles via sequencing total DNA from the cluster.

We found that a 54% of the total MCs was not able to grow on Mo(VI) at all concentrations while the rest of MCs (46%) were able to grow at least on 1 mM Mo(VI). To map only regions effectively implicated in the phenotypic variation of the evolved strain, the resistant MCs were grouped into two main clusters, indicated as Resistant-Parental and Resistant-Evolved, on the base of the phenotype showed (S1 Fig). In the first cluster we gathered all clones that did not grow on the media with 5 mM Mo(VI) and had white or light blue colonies on the media with 2.5 mM Mo(VI) like the UMCC 855 strain. In the second one we clustered all clones that, like UMCC 2581, had dark blue colonies on all the Mo(VI) media arranged (Table 2). The sensitive clones or the other ones that showed intermediate phenotypes or resistance were not considered. Successively, the genomic DNA of the two main clusters along with gDNA of the parental and evolved strains was extracted and sequenced (S1 Table).

**Table 2.**
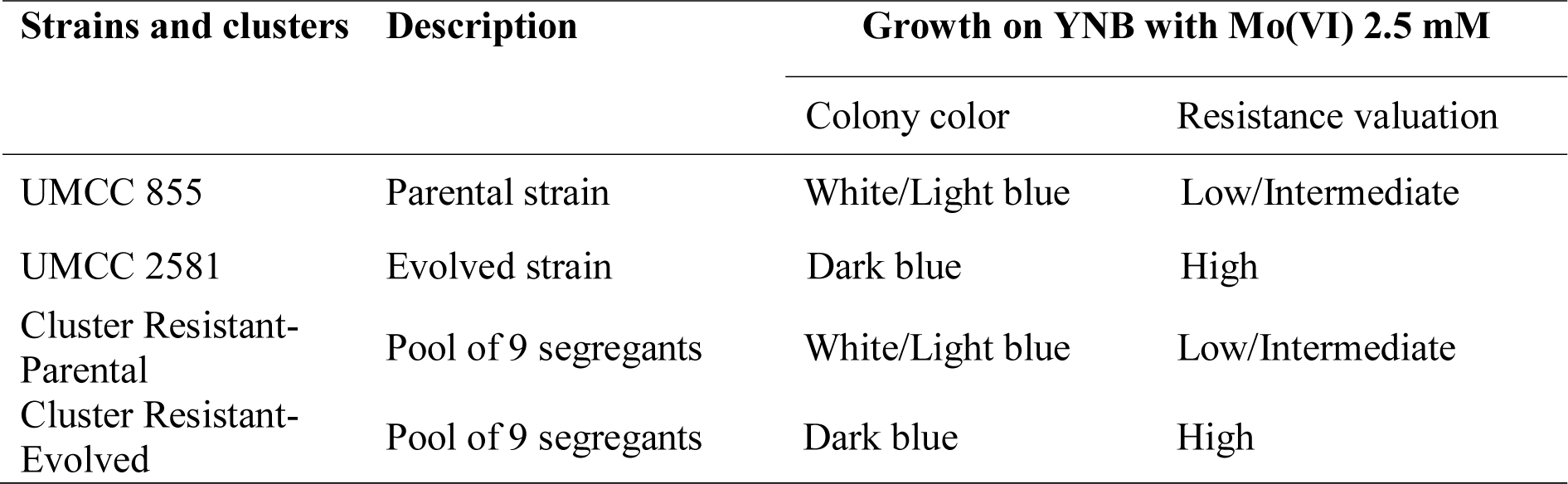
Clustering of the segregants obtained from the strain UMCC 855

| Strains and clusters | Description | Growth on YNB with Mo(VI) 2.5 mM |  |
| --- | --- | --- | --- |
|  |  | Colony color | Resistance valuation |
| UMCC 855 | Parental strain | White/Light blue | Low/Intermediate |
| UMCC 2581 | Evolved strain | Dark blue | High |
| Cluster Resistant-Parental | Pool of 9 segregants | White/Light blue | Low/Intermediate |
| Cluster Resistant-Evolved | Pool of 9 segregants | Dark blue | High |

The sequence reads were then aligned with the S288c reference sequence and only the sites heterozygous in the UMCC 855 parental strain were selected and used in the subsequent analysis. The allele frequency ratio p-value of the selected variants in the DNA of the clusters Resistant-Parental and Resistant-Evolved was then plotted against the variant position on the chromosome. The result is shown in Fig 2a. The cutoff threshold chosen at 20 log units allowed the identification of four QTLs: two peaks were present on chromosome 4, one on chromosome 6 and one on chromosome 12. The width of each peak was determined by dropping 5 log units from the top of the peak.

**Fig 2.**
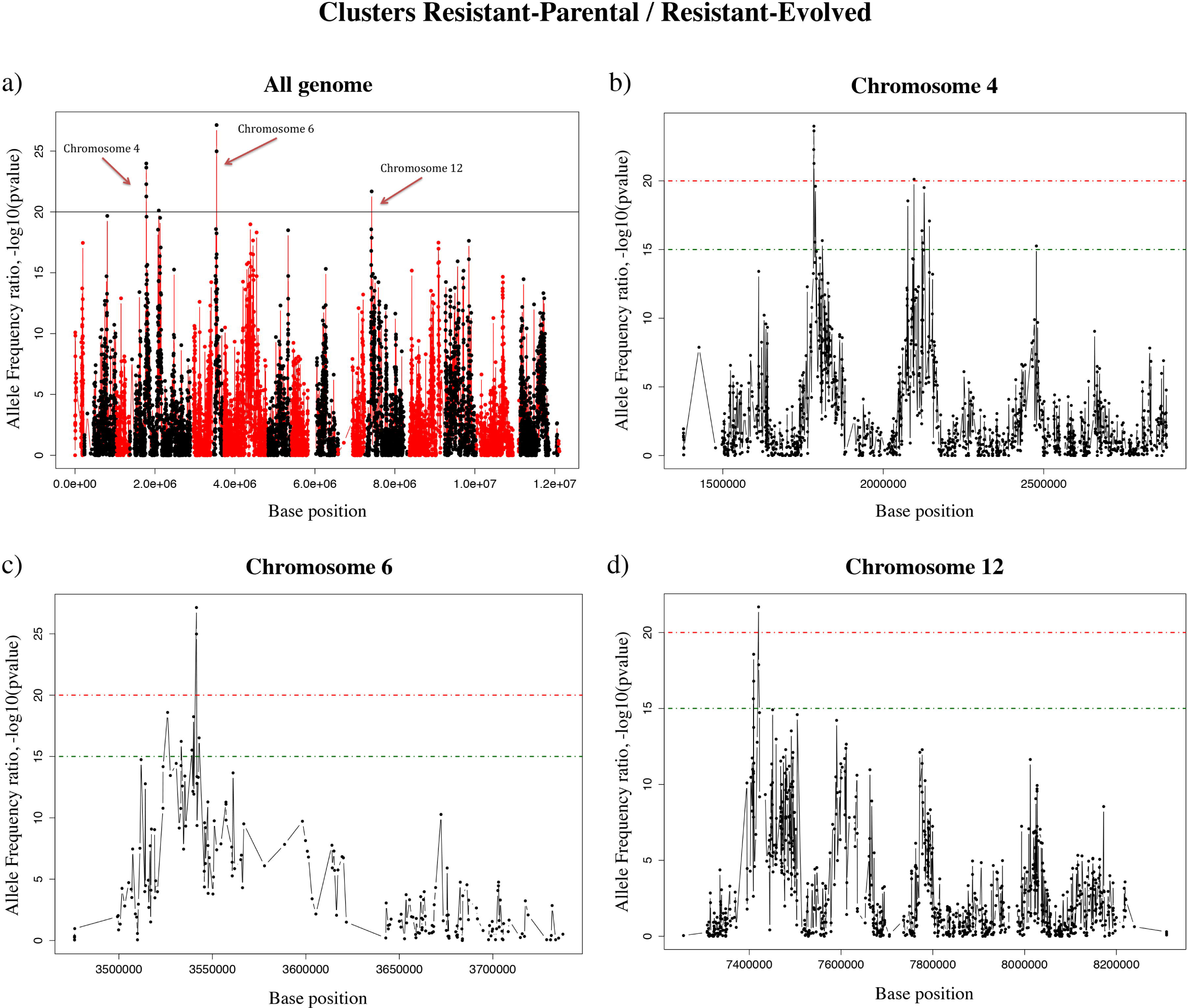
Allele frequency ratio plot. The variants −log 10 p-values for Fisher’s Exact Test (each represented by one dot) comparing clusters Resistant-Parental and Resistant-Evolved are shown. The QTL cutoff threshold was chosen at 20 log unit (black in a or red dashed line in b, c, d), the green dashed line represents the width of each peak (15 log unit), a) The dots colors (red and black) alternate when change the chromosome of reference, starting from the red chromosome 1 on the left. Candidate QTLs were found on chromosomes 4, 6 and 12. b) Allele frequency ratio plot for chromosome 4. c) Allele frequency ratio plot for chromosome 6. d) Allele frequency ratio plot for chromosome 12.

The QTLs present on chromosome 4 (Fig 2b) presented the larger peaks with a width of about 26 Kbp (between 423000 and 449000 bp) and 67 Kbp (between 717000 and 784000 bp) respectively for peak 1 and 2. The numbers of genes in these intervals were high, with 19 genes identified in peak 1 and 34 genes in peak 2 (S2 Table).

The peak on chromosome 6 (Fig 2c) showed a width of about 17 Kbp (between 57000 and 74000 bp) but the minimum number of genes (6) found (S3 Table).

The narrowest peak was found on chromosome 12 (Fig 2d) with a width of about 11 Kbp (between 164000 and 175000 bp) and 9 genes detected (S4 Table).

In order to identify the genes underlying the QTL, we initially examined the *S. cerevisiae* genome database (http://www.yeastgenome.org). Considering all peaks together, in the 121 Kbp mapped region 68 genes were annotated. Among these seven were dubious ORFs (YDL016C, YDL011C, YDR133C, YDR136C/VPS61, YDR149C, YDR154C and YDR157W) and seven were proteins of unknown function (YDL009C, YDL007C-A, YDL012C, YDR132C, YDR161W, YLR012C and YFL034W) that we did not consider any further. Besides dubious ORFs and unknown function protein, seven genes were ascribed to cellular process involved in reproduction *(CDC7, APC11, MCD1, RMD1, SWI5, CPR1* and *NSE1).* Two genes were attributed to fatty acid/phospholipid metabolism *(TSC13* and *EKI1)* and five genes to transport function *(ERP3, DOP1, PEX7, ENT5* and *SEC1).* Moreover, nine genes to transcription/translation process *(NOP1, MTQ2, TAF12, CTH1, GIR2, RPA14, CWC15, RPL22B* and *PPR1)* and three genes to mitochondrial function *(ATP16, KGD2* and *PAM18).* Finally, five genes to ubiquitin/proteasome process *(SLX5, RPT2, YDR131C, RUB1* and *SAN1)* and nine genes to cell integrity *(NHP10, FIN1, MKC7, NBP2, TUB2, MOB2, TEN1, GAT3* and *BRE2).* None of these genes had any obvious relationship to GSH production, resistance to metals or to oxidative stress. In contrast, *PTC1* and *NUM1* have roles in metals resistance mechanisms [35,36] and *SSY1* in regulation of GSH precursor amino acids permeases [37]. However, these genes did not show any genomic variations between the parental and evolved strain (on *NUM1* sequence there was reads alignment bias that eliminated any variant calls in the gene). On the other hand, *GRX6* and *MED2* on chromosome 4 peak 1, *YCF1, RGP1, HPR1, HOM2* and *SAC3* on chromosome 4 peak 2, *RPO41* and *RIM15* on chromosome 6 and *RLP24* and *LOT6* on chromosome 12 were considered to be worth for further investigation. In particular, comparing the UMCC 855 and UMCC 2581 coding regions, *GRX6* that encodes for a glutaredoxin involved in oxidative stress response, showed a single nucleotide polymorphism (SNP), whereas *MED2* gene, that encodes for a subunit of the RNA polymerase II mediator complex, involved as well in oxidative stress response, showed one SNP and two insertions (S2 Table). Instead, on chromosome 4 peak 2, *YCF1* (Yeast Cadmium Factor: vacuolar glutathione S-conjugate transporter), *RGP1* (gene implicated in retrograde transport from endosome to Golgi), *HPR1* (components of conserved THO nuclear complex) and SAC3 (mRNA export factor), were all related to metals detoxification. The *YCF1* gene differed in UMCC 2581 compared to UMCC 855 sequence, only for a single SNP in the coding region, two in case of *HPR1* and three in both, *RGP1* and *SAC3* (S2 Table). *HOM2*, the last gene annotated on chromosome 4 peak 2, which catalyzes the second step for methionine biosynthesis, displayed a single variant in the genetic sequence. Regarding the genes annotated on peaks on chromosomes 6 and 12, they were all related to oxidative stress response. On chromosome 6, the mitochondrial RNA polymerase RPO41 showed one SNP in the coding region. The protein kinase *RIM15* showed the higher number of genomic variations in a single gene with ten SNPs and one insertion (S3 Table). Finally, both genes annotated on chromosome 12, displayed one SNP comparing parental and evolved DNA sequences (S4 Table).

### Identification of new mutations

In order to identify new mutations, we performed the sequence analysis of gDNA from both parental and selected evolved strain (S1 Table). To catalogue copy-number variation (CNV) in the UMCC 855 and 2581 yeast genomes, the depth of sequencing coverage for each genome was calculated (Fig 3). While the parental strain exhibited a normal chromosomal set (Fig 3a), the evolved strain revealed a whole-chromosome amplification (Fig. 3b). The read depth of chromosome 1 was 1.5-fold greater than the median of the strain, pointing out the presence of an extra-copy of this chromosome in UMCC 2581.

**Fig 3.**
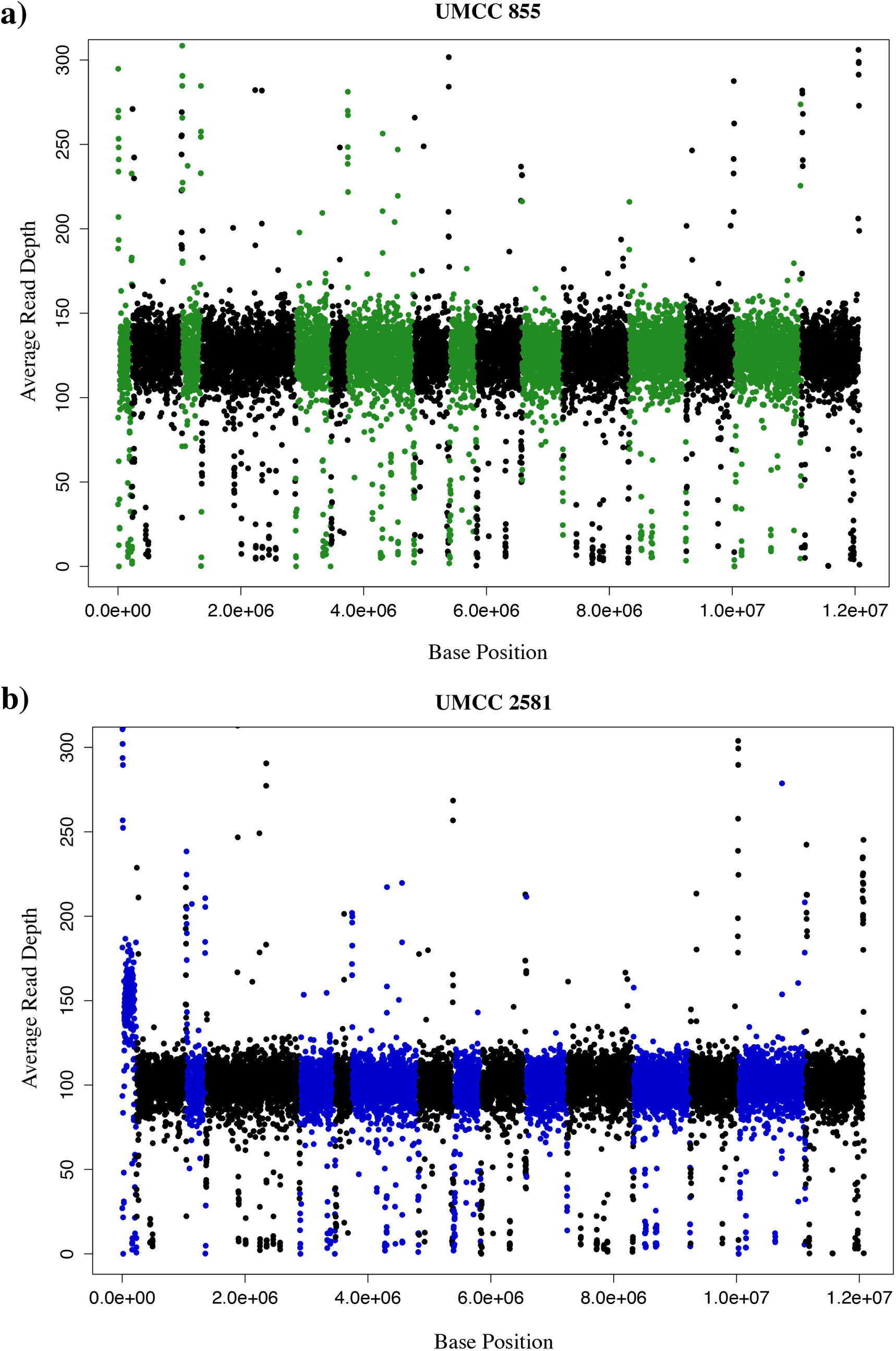
Chromosomal aneuploidy determined by whole-genome sequencing coverage. The average sequencing coverage was determined using a sliding window of 1000 bp and plotted in chromosomal order (each represented by a colored dot). The dot colors (green or blue and black) alternate between chromosomes of the reference, starting from the green or blue chromosome 1 on the left, a) Coverage plot of UMCC 855 parental strain, b) Coverage plot of UMCC 2581 evolved strain.

The UMCC 855 and UMCC 2581 sequences of the genes within the QTLs regions were compared to find any new mutations present in the evolved but not in the parental strain. Among all genes evaluated only two genes, *MED2* and *RIM15* both related to oxidative stress response, presented new mutations in the UMCC 2581 strain sequence and were potentially related to the evolved phenotype. In particular, the *MED2* sequence presented an inframe insertion C > CTGTTGTTGT in ORF position 441267 (S2 Table) that was not present in UMCC 855 sequence, while a new mutation observed in the *RIM15* sequence was a frameshift mutation (GA > G) in position 1066 (S3 Table).

### Transcriptome profile comparison of parental and evolved strains

For a comprehensive evaluation of the occurred genome alteration underlying the higher GSH production and resistance to Mo(VI), we assessed gene expression levels of the evolved and parental strains. Gene expression was measured at three quarters of the way through the exponential phase. This point was chosen in order to precede the major transcriptional reprogramming event during fermentation of a synthetic must that is triggered during entry into stationary phase [21,38]. Under the chosen condition, the transcriptome is stable and expected to provide a relevant picture regarding the different strains capacity to produce GSH in mimicking natural must condition.

The transcriptomes of the UMCC 2581 evolved strain and the UMCC 855 parental strain were first compared in the expression profile plot (Fig 4). The graph shows a higher average expression of the genes present in the chromosome 1 of the evolved strains 2581 (0.57) comparing to the average expression of all other chromosomes taking together. Since the average expression in a comparison between two strains with a normal chromosomal asset is expected to be 0, this result confirm, as previously described (Fig 3), the presence of an extra copy of chromosome 1 in UMCC 2581 strain. Accordingly with the demonstrated aneuploidy, all UMCC 2581 differentially expressed genes present on chromosome 1 (47 genes) were not considered in the subsequent analysis. Without counting the chromosome 1, we observed that 296 genes were differentially expressed between the two strains at an FDR < 0.05 (Fig 5 and S5 Table). Of the 161 genes up-regulated, 66 genes modified their expression more than two-fold as well as 61 genes out of 135 genes down-regulated. Our results indicate that a small fraction of the genes in the genome are differentially expressed but with quite a large subset (almost half) displaying strong variations.

**Fig 4.**
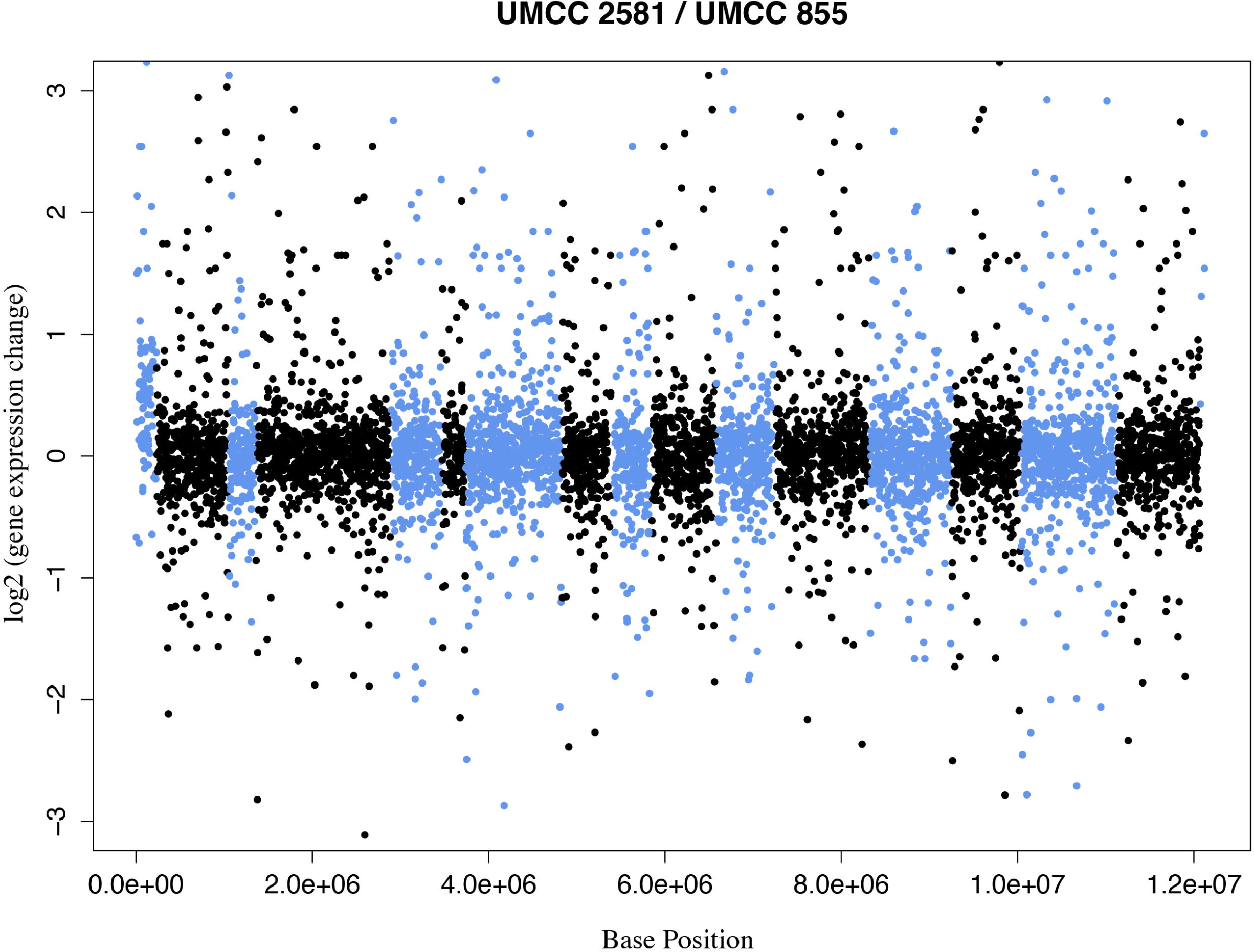
Expression profile plot comparing the UMCC 2581 to UMCC 855. Each dot represents the log2 of the gene expression fold-change comparing the two strains. The dots colors (light blue and black) alternate when change the chromosome of reference, starting from the light blue chromosome 1 on the left.

**Fig 5.**
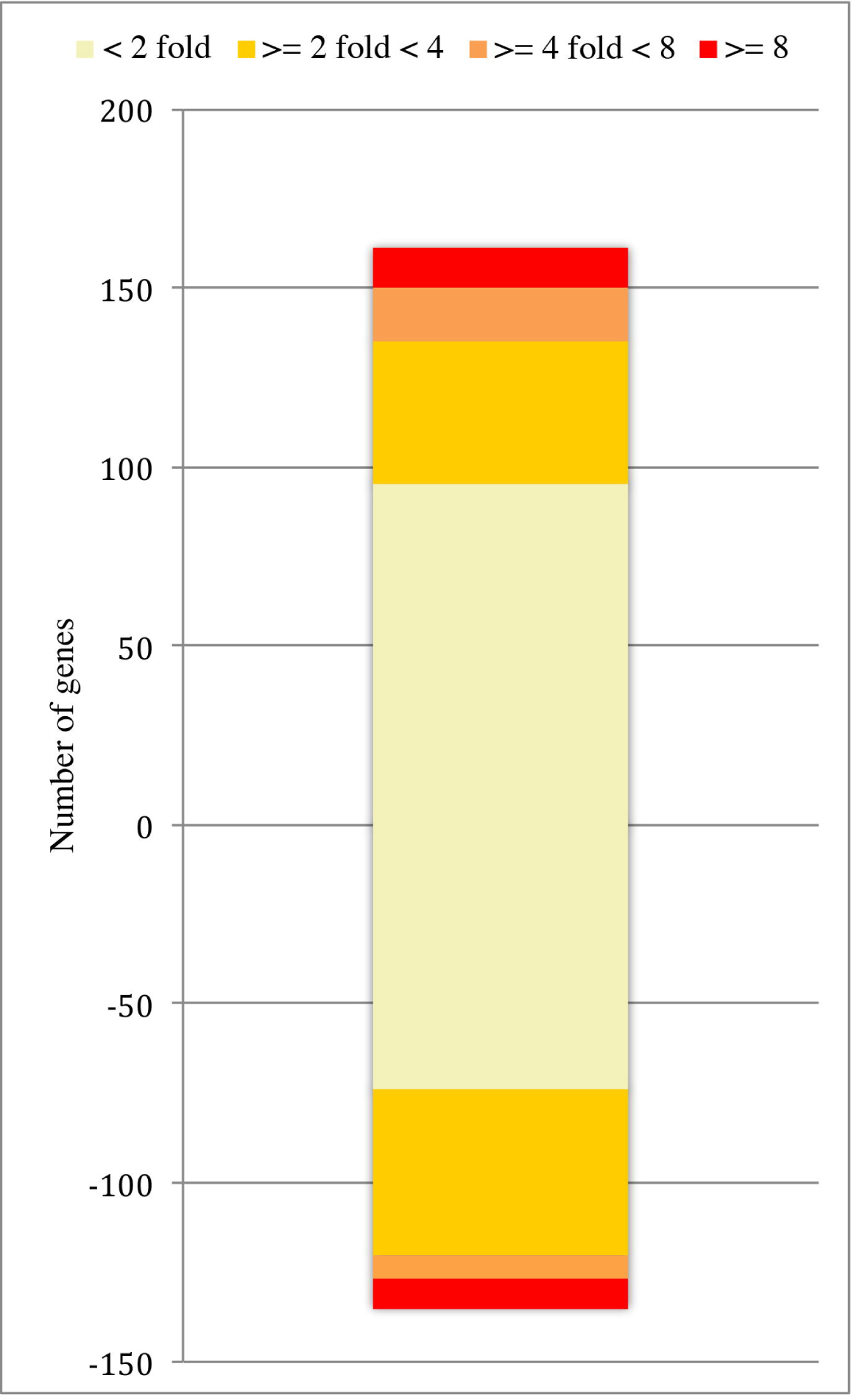
Genes differentially expressed in the comparison between the strains UMCC 855 and UMCC 2581. The positive values of the vertical axis represent the number of genes more expressed in UMCC 2581 in comparison to UMCC 855, while the negative values reports the number of genes down expressed. Four different thresholds were considered, the genes with a low expression difference (light green and orange, differentially expressed between 2- and 4-fold), those with a medium expression difference (dark orange, between 4- and 8-fold) and those with high expression difference (red, more than 8-fold). FDR<0.05.

Among the top 10% of genes that were strongly over-expressed (16 genes, S5 Table), UMCC 2581 exhibited a small but significant set of permease genes including two amino acids permeases *(DIP5* and *GNP1)* involved in transport of GSH precursor amino acids (cysteine, methionine, glutamate and glycine) and *SUL1* gene involved in sulfate assimilation pathway. On the other hand, the top 10% of genes strongly under-expressed (14 genes), was characterized by the null production of two genes *(MAL11* and *MAL13)* that could impair the strain ability to utilize maltose.

Regarding the genes mapped into the QTL regions, the vacuolar GSH S-conjugate transporter, *YCF1,* was the only differentially expressed gene detected comparing UMCC 2581 to UMCC 855 transcriptomes. This important gene in the resistance to heavy metals, showed a slightly overexpression in UMCC 2581.

### Gene Ontology (GO) analysis

To have a more accurate comparison between strains, enriched functional classes of genes were obtained using “GO Term Finder” within the SGD database. GO term annotation analysis was used to detect enriched biology process (S6 Table), function (Table 3 and S7 Table) and component (S8 Table) in UMCC 2581 compared to UMCC 855.

**Table 3.**
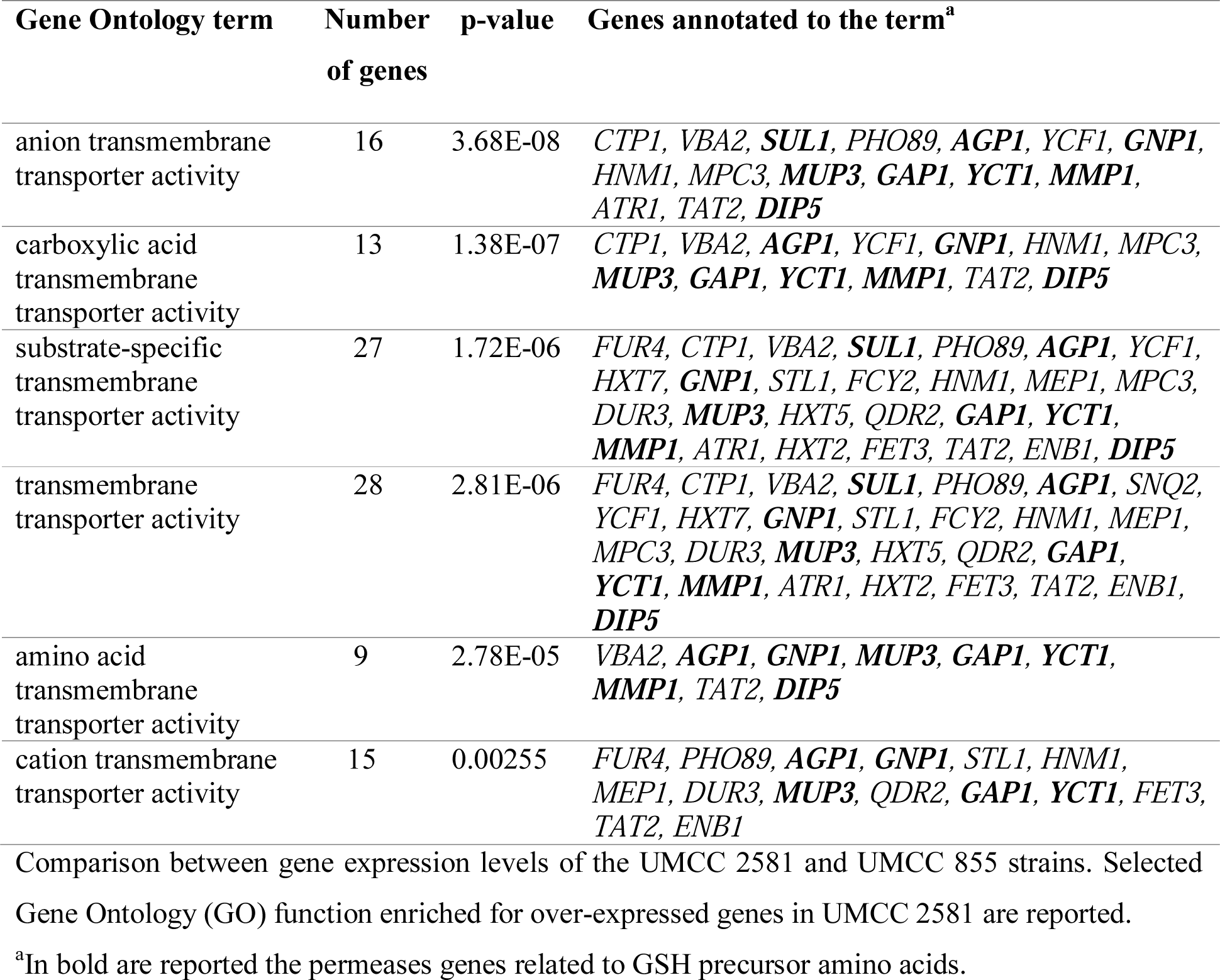
Selected Gene Ontology - Function.

All the GO analysis performed displayed consistent results that revealed, for the up-regulated genes, the term related to the transport activity striking enriched. Conversely, the same analysis considering the down-regulated genes, exhibited enriched terms for only for four genes *(THI2, THI4, THI20, THI21)* related to metabolism of thiamine in GO term “Process”. The graphical analysis of the GO term “molecular function” (Fig 6), showed that within the dataset of transporter activity, the terms associated with the amino acids transporter were particularly enriched. Coherently, as observed in Table 3, many of the genes that strongly characterize all the enriched function classes, were genes involved in amino acids transporter. Noteworthy, 7 out of 9 genes in the GO class “amino acid transmembrane transporter activity” (genes in bold in Table 3) were genes related to GSH precursor amino acids. In particular, Dip5p (more than 20 fold-change) mediates high-affinity transport of L-glutamate but it is also a transporter for glycine, *YCT1* and *MUP3* encode, respectively, for high-affinity cysteine transporter and low affinity methionine permease, Gnp1p transports both as well as Agp1p together with glycine and Gap1p is a general amino acid permease. Finally, even though not a transporter for amino acid, the high affinity sulfate permease, encoded by *SUL1,* is involved in methionine and cysteine biosynthetic process by the sulfate assimilation pathway.

**Fig 6.**
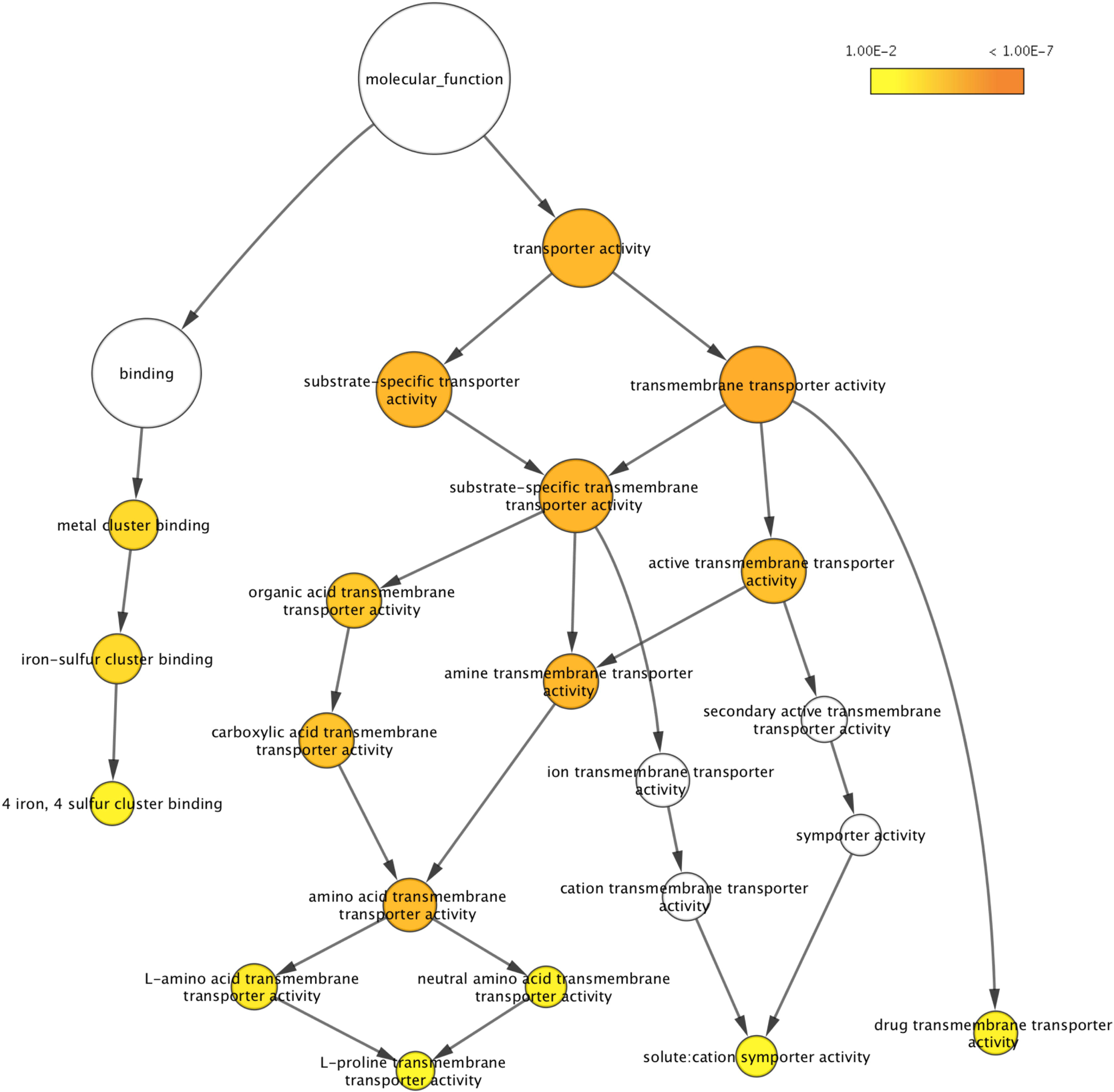
BiNGO result for the over-expressed genes in UMCC 2581 considering the GO term “Molecular function”. The node size corresponds to the number of proteins that are assigned with the individual term. The terms with a p-value below 0.01 were defined as significant (yellow), and a darker color represents a lower p-value (orange < 1.00E-7). White nodes are not significantly over-represented; they are included to show the colored nodes in the context of the GO hierarchy.

Furthermore, although not present in enriched GO classes, also *MET5, GLT1* and *SER3* genes were found over-expressed in UMCC 2581. These genes were involved in biosynthesis of methionine and cysteine (*MET5*, sulfite reductase beta subunit), glutamate from glutamine and alpha-ketoglutarate *(GLT1,* NAD(+)-dependent glutamate synthase) and glycine *(SER3,* 3-phosphoglycerate dehydrogenase).

## Discussion

The relationship between metal uptake and toxicity has been documented in many instances since metal resistant microbial strains often exhibit an ability to prevent or reduce entry of toxic metal species into the cell [39,40].

Among the events conferring resistance to sulfate toxic analogues, a mutation in the high-affinity permeases encoded by the genes *SUL1* and *SUL2* is one of the most probable [41–43]. In this case, the subsequent impaired assimilation of SO2 can result in a low or nil sulphites and sulphides production [11]. Another resistance mechanism of the cells could be related to the production of GSH, which is known to have an essential role in the defense against oxidative stress and metal toxicity [43,44]. In particular, GSH biosynthesis in *S. cerevisiae* takes place in two ATP-dependent steps. In the first, cysteine is linked with glutamate by γ-glutamylcysteine synthetase (encoded by *GSH1)* to form γ-glutamylcysteine. In the second step, glycine is added to this intermediate product by glutathione synthetase (encoded by *GSH2)* to form the final product [45,46]. Several authors have reported that GSH is able to chelate heavy metals by forming complexes (metal-GSH complex) that are actively transported into the vacuole or removed from the cell by specific transporters such as Ycf1p and Gex1p [12,47–50].

### Chromosomal variations in the evolved strain

*Saccharomyces cerevisiae* wine yeasts are characterized by the complexity of their nuclear genome and, rather than being strictly diploid, many strains display chromosomal copy number variation (polyploidy, aneuploidy) or rearranged chromosomes [51–53]. In this work, we observed that the median read depth of UMCC 2581 chromosome 1 was greater by 1.5-fold (3:2 ratio represent at least one extra genomic copy in a diploid strain) than the median of the strain. This pointed out the presence of an extra copy of this chromosome (Fig 3b) and confirming the high level of polymorphism in wine yeasts. Accordingly, this large-scale genomic reorganization provided an average high expression of all genes present in chromosome 1 of the evolved strains UMCC 2581 (117 ORF, Fig 4). Moreover, the chromosome 1 aneuploidy besides the increased expression of the involved genes, raises also the possibility of both new mutations and different segregation of parental alleles. However, how these modifications can affect the different phenotype is unknown and hard to define.

### Identification of candidate genes in QTL

Through the analysis of the two resistant clusters of interest (Resistant-Parental/Resistant-Evolved), obtained with our approach, it was possible to map the QTL that characterize the evolved phenotype. The allele frequency plot revealed four major loci on chromosomes 4, 6 and 12 where 68 genes were annotated. Among these, eleven genes (*GRX6, MED2, YCF1, RGP1, HPR1, HOM2, SAC3, RPO41, RIM15, RLP24* and *LOT6*) presented genomic variations comparing parental and evolved strains sequences and were functional related to GSH production, resistance to metals or to oxidative stress.

The *HOM2* gene is the only candidate gene that could be related to GSH production by its amino acid precursor biosynthesis. Indeed, the gene product, aspartic β-semialdehyde dehydrogenase, catalyzes the second step of the threonine and methionine metabolic pathway starting from aspartic acid [54]. The *HOM2* sequence displayed only one single nucleotide polymorphism, but it occurs in the NAD binding domain. Therefore, we can speculate that the resulting replacement of histidine by asparagine (His29Asn, S2 Table), both uncharged amino acids but with different 3D-structure, could affect the efficiency of binding NAD and, consequently, the catalytic efficiency of the protein.

On behalf of its function and its close relation with GSH, the most convincing candidate gene in relation with the metals resistant phenotype, seemed to be the Yeast Cadmium Factor *(YCF1),* which encodes for a well-studied ATP-binding cassette (ABC) protein localized in vacuolar membrane as glutathione S-conjugate (GS-X) transporter [47]. Szczypka and co-workers [55], who discovered *YCF1,* described its ability to confer cadmium resistance. Indeed, the Ycf1p is also able to transport into the vacuole a broad range of heavy metals as well as xenobiotic substrates providing resistance to cells [48,49,56-58]. The SNP found as heterozygous in UMCC 855 sequence, and fixed as homozygous alternative allele in UMCC 2581, changed the glutamine amino acid in position 899 with a histidine (S2 Table). This variation occurred in the regulatory domain and in particular next to K890, an ubiquitination site [59,60]. No DNA variations were exhibited on the consensus element YRE (binds by Yap1p, the transcriptional activator) [61], moreover, RNA-seq experiments showed a slight increased *YCF1* expression in UMCC 2581. Taking into account these findings, we have hypothesized that the presence of His, possibly for steric hindrance, reduced or prevent the ubiquitination of lysine 890 leading to a lower degradation of the protein.

Although the vacuole emerges as a major hot-spot for metal detoxification, a number of pathways that play a more general, less direct role in promoting cell survival under stress conditions as mRNA processing and transport, can be identified. In this context, on chromosome 4 peak 2 besides Ycf1p, also the product of *RGP1, HPR1* and *SAC3* genes were found to be involved in the metal detoxification (S2 Table).

Metal toxicity may be caused by impaired DNA repair, inhibition or disturbing of enzyme function but also by oxidative stress that originates from toxic levels of oxygen-derived reactive species (ROS) stimulated directly or indirectly by metals [58,62,63]. ROS attack and damage all cellular macromolecules, leading to protein oxidation, lipid peroxidation and DNA damage [43]. For this reason, proteins related to oxidative stress were considered as candidate genes in our analysis and they were the major represented function (6 out of 11 genes). Among these, Grx6p, Rim15p and Lot6p have proven to be directly involved in the oxidative stress response [64,65]. Moreover, the *RIM15* gene, that presented a new mutation compared to the parental strain, provided the most interesting sequence modification. Rim15p is a protein kinase by which *Saccharomyces cerevisiae* regulates the post diauxic shift, entry into meiosis and stationary phase and life-span [66,67]. The reduced life-span observed in rim15 deletion cells was probably due to their deficiency in oxidative damage prevention [68], indeed the Rim15p regulon comprises genes implicated in oxidative stress. In our observation, the *RIM15* gene was a hot-spot of mutations, gathering eleven polymorphism between the two sequences. Noteworthy, none of them were in the protein kinase domain even though a frameshift mutation (GA > G) in position 1066 (S3 Table) observed in one allele of evolved strain, arose between the two protein kinase domain, probably decreasing or deleting the protein functionality.

As mentioned before regarding metals detoxification, some housekeeping processes appear to play a significant role also in case of hyperoxia resistance. Consistent with these observations, in our QTL genes involved in controlling the activity of general transcription factors and RNA polymerase II [69,70] as Med2p, in the production of 60S ribosomal subunits [71] as Rlp24p, or encoding for a mitochondrial RNA (mtRNA) polymerase [72] as Rpo41p were found. The specific function carried out in response to oxidative stress by these genes is not clearly understood, however, their involvement in the response against oxidative stress was reported by several authors [73–76].

The deeper analysis of the sequences revealed that in four genes, *HPR1* and *HOM2* on chromosome 4 and the two candidate genes on chromosome 12 *RLP24* and *LOT6,* the UMCC 2581 evolved strain presented the restored reference allele. This suggests that in these cases the alternative alleles arose in parental strain leading to a loss rather than a gain of function. The corresponding fully functional proteins, in particular Hpr1p and Lot6p where the alternative alleles brought a frameshift and a stop codon UMCC 855, probably provide a contribution in the resistant phenotype. On the contrary, in some other cases the mutations observed as heterozygous in UMCC 855 and fixed in the evolved strain, as in *MED2,* or new mutations in the evolved strain, as in *RIM15,* seems to result in a corrupted protein. This unexpected protein dysfunction, reported only in UMCC 2581, could nevertheless comply with the mechanisms of GSH overproduction recently proposed by Zhu et al. [77]. In this work, the authors proposed that, first mutant cells accumulated high ROS levels because of deficient mutated protein, and then the accumulated endogenous ROS subsequently led to chronic oxidative stress and triggered the oxidative stress response, resulting in overproduction of GSH. However, the suggested hypothesis and the mutations' effects proposed requires further study to be confirmed.

### Transcriptome profiles comparison

To find relevant evidence supporting the different strains capacity to produce GSH, UMCC 855/2581 genes expression levels were assessed. The SM medium was used in order to mimicking natural must condition and the sampling point was chosen to avoid the most critical phase at the beginning of the fermentation and the major transcriptional reprogramming event that triggered entering into the stationary phase. The most abundant overexpressed GO classes in UMCC 2581 were all involved in transport activity strongly underlying how the major differences between the two strains were situated in this process. Important to notice, the terms associated with the amino acids transporter were particularly extended in the graphical representation (Fig 6), but also highly characterizing the other enriched function classes (Table 3). A deeper analysis of the gene annotated in ‘amino acid transmembrane transporter activity’ revealed, remarkably, that 7 out of 9 were genes related to all GSH precursor amino acids. Cysteine and methionine are transported by Yct1p, Mup3p, Gnp1p and Agp1p, glutamate and glycine by Dip5p (the most differentially higher expressed gene) and Agp1, all are transported by Gap1p, a general amino acid permease [78–80]. Moreover, among the differentially over-expressed genes, other genes potentially related to GSH production were found (S5 Table). In particular *SUL1,* present together with the above-mentioned amino acids permeases in numerous GO enriched classes, encodes for high affinity sulfate permease [81]. *SUL1* along with *MET5,* that encodes for the β-subunit of the *S.cerevisiae* sulfite reductase [82], are involved in the sulfate assimilation pathway that precede the synthesis of sulfur-containing amino acids cysteine and methionine [83]. Another gene is *GLT1,* which encodes for GOGAT (glutamate synthase) and synthesizes two molecules of glutamate out of one molecule of glutamine and one molecule of α-ketoglutarate [84]. Finally, Ser3p, phosphoglycerate dehydrogenase, catalyzes the first reaction of serine and glycine biosynthesis from the glycolytic metabolite 3-phosphoglycerate [85]. Therefore, our results evidenced that all the GSH precursor amino acids (sulfur-containing amino acids, glutamate and glycine) were over-expressed in its biosynthetic pathway, from permeases to synthetic enzymes (Fig 7). Thus, it is suggested that this aptitude to collect the precursor amino acids from the media, considering also the over-represented biosynthetic steps in each amino acid pathway, might lead to an overproduction of glutathione by providing large amount of precursors. The gene expression pattern here observed is of particular importance for the technological point of view: although genomic regulation may differ in natural musts with a different nutritional status, the ability to gather precursor amino acids from the media is probably relevant for a constantly high GSH production in real fermentations where must are different year by year.

**Fig 7.**
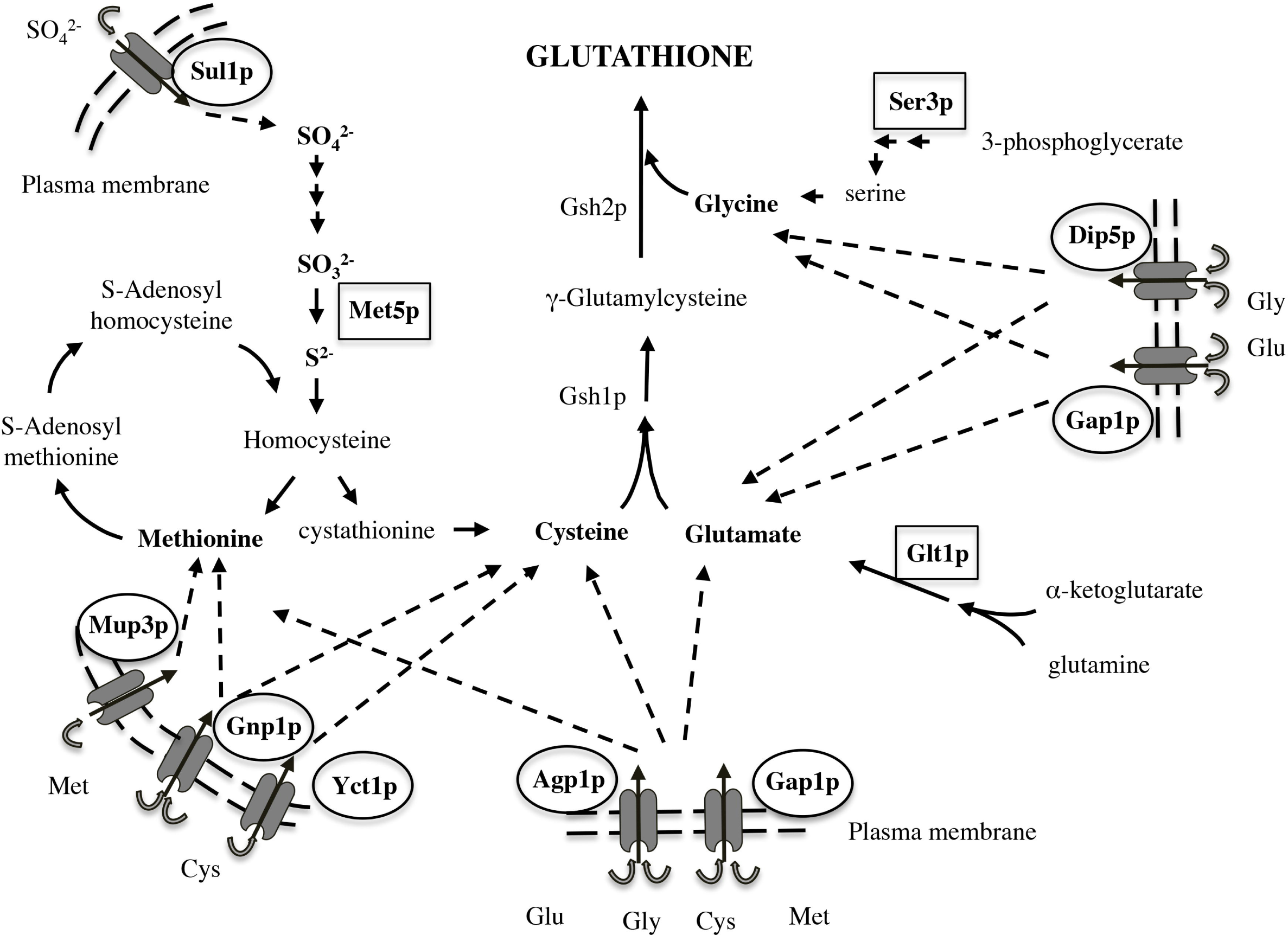
Glutathione precursor amino acids, permeases and enzymes, involved in the hypothesized machanism for the high GSH production.

The analysis of genes related to GSH metabolism allowed us to detect two over-expressed GSH-related genes that are involved in metals detoxification: *YCF1* and *ECM38*. The *YCF1* gene was the only one detected in both analyses, whole-genome and transcriptome sequencing, and its important role in providing resistance to heavy metals and xenobiotics as GSH S-conjugate transporter was previously described. The Y-glutamyltranspeptidase (Y-GT), in *S. cerevisiae* encoding by *ECM38*, is the major GSH-degrading enzyme. Once it is transported into the vacuole, GSH is degraded by the vacuolar membrane-bound Y-GT and L-cysteinyl glycine dipeptidase by the cleavage of the Y-glutamyl moiety and the release of cysteinylglycine, further degraded to its constitutive amino acids [86]. Analogue mechanism might be responsible for the recycle of xenobiotics/metal-GSH complex stored in the vacuole, which can be excreted from cells [87,88] suggesting a possible mechanism for molybdate resistance in evolved strain.

To outline, the analysis of the transcriptional profiles revealed two very important aspects. The first is the global over-expression of the amino acids permeases, noteworthy especially for the final purpose of the evolved strain: high GSH production in oenological applications. The second aspect is the remarkable role of transport processes in the definition of the desired phenotype. This was evident in transcriptome analysis, where was the only enriched process, and also in the QTL analysis, where we detected key genes related to transport activity. Regarding the application of molybdate as selective pressure to obtain evolved strains, *YCF1,* detected in both analyses, emerged as a major hot spot for metal detoxification. Besides the *YCF1,* a number of genes associated with a more general transport pathways (for example vesicle and nucleocytoplasmic transport), probably play also a role in promoting cell survival under metal/oxidative stress conditions and in the GSH production and homeostasis.

## Conclusion

In this work, we applied quantitative genetics to study the genetic changes underlying the high GSH production showed by the wine *S. cerevisiae* strains UMCC 2581 selected in a molybdate-enriched environment after sexual recombination.

We identified four peaks within 11 candidate genes in QTL analysis and 296 genes differentially expressed between parental and evolved strain. The complex genetic traits and the wide variations produced by sexual recombination resulted in a presumed additive phenotype effects.

The high GSH production phenotype included over-expression of amino acids permeases and precursor biosynthetic enzymes rather than the two GSH metabolic enzymes, whereas GSH production and metabolism, transporter activity, vacuolar detoxification and oxidative stress response enzymes were probably added resulting in the molybdate resistance phenotype.

A thorough understanding of the genes variations effects and the scope of the aneuploidy consequence on chromosome 1 are necessary to address the exact relationships between the evolved phenotypes and candidate genes expression.

This study provides an example of a combination of an evolution-based strategy to successful obtain yeast strain with desired phenotype and inverse engineering approach to genetic characterize the evolved strain.

The genetic information provided could be useful for further optimization of the evolved strains and for providing an even more rapid approach to identify new strains, with a high GSH production, through a marked-assisted selection strategy.

## References

1. Sonderegger M, Sauer U. Evolutionary Engineering of *Saccharomyces cerevisiae* for Anaerobic Growth on Xylose. Appl Environ Microbiol. 2003;69(4): 1990–8.

2. Kuyper M, Toirkens MJ, Diderich JA, Winkler AA, van Dijken JP, Pronk JT. Evolutionary engineering of mixed-sugar utilization by a xylose-fermenting *Saccharomyces cerevisiae* strain. FEMS Yeast Res. 2005;5(10): 925–34.

3. Stanley D, Fraser S, Chambers PJ, Rogers P, Stanley GA. Generation and characterisation of stable ethanol-tolerant mutants of *Saccharomyces cerevisiae*. J Ind Microbiol Biotechnol. 2010;37(2): 139–49.

4. Cadière A, Ortiz-Julien A, Camarasa C, Dequin S. Evolutionary engineered *Saccharomyces cerevisiae* wine yeast strains with increased in vivo flux through the pentose phosphate pathway. Metab Eng. 2011;13(3): 263–71.

5. Kutyna DR, Varela C, Stanley GA, Borneman AR, Henschke PA, Chambers PJ. Adaptive evolution of *Saccharomyces cerevisiae* to generate strains with enhanced glycerol production. Appl Microbiol Biotechnol. 2012;93(3): 1175–84.

6. Sipiczki M. Diversity, variability and fast adaptive evolution of the wine yeast (*Saccharomyces cerevisiae*) genome-a review. Ann Microbiol. 2011;61(1): 85–93.

7. Cakar ZP, Turanli-Yildiz B, Alkim C, Yilmaz U. Evolutionary engineering of *Saccharomyces cerevisiae* for improved industrially important properties. FEMS Yeast Res. 2012;12(2): 171–82.

8. Winkler JD, Kao KC. Recent advances in the evolutionary engineering of industrial biocatalysts. Genomics. 2014;104(6): 406–11.

9. Santos CNS, Stephanopoulos G. Combinatorial engineering of microbes for optimizing cellular phenotype. Curr Opin Chem Biol. 2008;12(2): 168–76.

10. Parekh S, Vinci VA, Strobel RJ. Improvement of microbial strains and fermentation processes. Appl Microbiol Biotechnol. 2000;54(3): 287–301.

11. De Vero L, Solieri L, Giudici P. Evolution-based strategy to generate non-genetically modified organisms *Saccharomyces cerevisiae* strains impaired in sulfate assimilation pathway. Lett Appl Microbiol. 2011;53(5): 572–5.

12. Mezzetti F, De Vero L, Giudici P. Evolved *Saccharomyces cerevisiae* wine strains with enhanced glutathione production obtained by an evolution-based strategy. FEMS Yeast Res. 2014;14(6): 977–87.

13. Marullo P, Bely M, Masneuf-Pomarede I, Aigle M, Dubourdieu D. Inheritable nature of enological quantitative traits is demonstrated by meiotic segregation of industrial wine yeast strains. FEMS Yeast Res. 2004;4(7): 711–9.

14. Giudici P, Solieri L, Pulvirenti A, Cassanelli S. Strategies and perspectives for genetic improvement of wine yeasts. Appl Microbiol Biotechnol. 2005;66(6): 622–8.

15. Hosiner D, Gerber S, Lichtenberg-Fraté H, Glaser W, Schüller C, Klipp E. Impact of acute metal stress in Saccharomyces cerevisiae. PLoS One. 2014;9(1): 1–14.

16. Pretorius IS. Tailoring wine yeast for the new millennium: novel approaches to the ancient art of winemaking. Yeast. 2000;16(8): 675–729.

17. Fay JC, McCullough HL, Sniegowski PD, Eisen MB. Population genetic variation in gene expression is associated with phenotypic variation in Saccharomyces cerevisiae. Genome Biol. 2004;5(4): R26.

18. Jackson P, Schacherer J. Population genomics of yeasts: towards a comprehensive view across a broad evolutionary scale. Yeast. 2016;33(3): 73–81.

19. Cromie GA, Hyma KE, Ludlow CL, Garmendia-Torres C, Gilbert TL, May P, et al. Genomic sequence diversity and population structure of *Saccharomyces cerevisiae* assessed by RAD-seq. G3. 2013;3(December): 2163–71.

20. Nagalakshmi U, Wang Z, Waern K, Shou C, Raha D, Gerstein M, et al. The transcriptional landscape of the yeast genome defined by RNA sequencing. Science. 2008;320(5881): 1344–9.

21. Treu L, Campanaro S, Nadai C, Toniolo C, Nardi T, Giacomini A, et al. Oxidative stress response and nitrogen utilization are strongly variable in Saccharomyces cerevisiae wine strains with different fermentation performances. Appl Microbiol Biotechnol. 2014;98(9): 4119–35.

22. Gobbi M, De Vero L, Solieri L, Comitini F, Oro L, Giudici P, et al. Fermentative aptitude of non-*Saccharomyces* wine yeast for reduction in the ethanol content in wine. Eur Food Res Technol. 2014;239(1): 41–8.

23. Bonciani T, Solieri L, De Vero L, Giudici P. Improved wine yeasts by direct mating and selection under stressful fermentative conditions. Eur Food Res Technol. 2016;242(6): 899–910.

24. Cherry JM, Hong EL, Amundsen C, Balakrishnan R, Binkley G, Chan ET, et al. Saccharomyces Genome Database: the genomics resource of budding yeast. Nucleic Acids Res. 2012;40(D1): D700–5.

25. Li H, Durbin R. Fast and accurate short read alignment with Burrows-Wheeler transform. Bioinformatics. 2009;25(14): 1754–60.

26. DePristo MA, Banks E, Poplin R, Garimella K V, Maguire JR, Hartl C, et al. A framework for variation discovery and genotyping using next-generation DNA sequencing data. Nat Genet. 2011;43(5): 491–8.

27. Cingolani P, Platts A, Wang LL, Coon M, Nguyen T, Wang L, et al. A program for annotating and predicting the effects of single nucleotide polymorphisms, SnpEff: SNPs in the genome of *Drosophila melanogaster* strain w 1118; iso-2; iso-3. Fly. 2012;6(2): 80–92.

28. Thorvaldsdottir H, Robinson JT, Mesirov JP. Integrative Genomics Viewer (IGV): high-performance genomics data visualization and exploration. Brief Bioinform. 2013;14(2): 178–92.

29. Giudici P, Kunkee RE. The effect of nitrogen deficiency and sulfur-containing amino acids on the reduction of sulfate to hydrogen sulfide by wine yeasts. Am J Enol Vitic. 1994;45(1): 107–12.

30. Ausubel FM, Brent R, Kingstone RE, Moore DD, Seidman JG, Smith JA, et al. Short Protocols in Molecular Biology, Volume 2. 5th ed. Wiley (USA); 2002. 1512 p.

31. Langmead B, Salzberg SL. Fast gapped-read alignment with Bowtie 2. Nat Methods. 2012;9(4): 357–9.

32. Anders S, Huber W. Differential expression analysis for sequence count data. Genome Biol. 2010;11(10): R106.

33. Maere S, Heymans K, Kuiper M. BiNGO: a Cytoscape plugin to assess overrepresentation of Gene Ontology categories in Biological Networks. Bioinformatics. 2005;21(16): 3448–9.

34. Parts L, Cubillos FA, Warringer J, Jain K, Salinas F, Bumpstead SJ, et al. Revealing the genetic structure of a trait by sequencing a population under selection. Genome Res. 2011;21(7): 1131–8.

35. Ruiz A, González A, García-Salcedo R, Ramos J, Ariño J. Role of protein phosphatases 2C on tolerance to lithium toxicity in the yeast *Saccharomyces cerevisiae*. Mol Microbiol. 2006;62(1): 263–77.

36. Paumi CM, Menendez J, Arnoldo A, Engels K, Iyer KR, Thaminy S, et al. Mapping Protein-Protein Interactions for the Yeast ABC Transporter Ycf1p by Integrated Split-Ubiquitin Membrane Yeast Two-Hybrid Analysis. Mol Cell. 2007;26(1): 15–25.

37. Forsberg H, Gilstring CF, Zargari A, Martínez P, Ljungdahl PO. The role of the yeast plasma membrane SPS nutrient sensor in the metabolic response to extracellular amino acids. Mol Microbiol. 2001;42(1): 215–28.

38. Rossignol T, Dulau L, Julien A, Blondin B. Genome-wide monitoring of wine yeast gene expression during alcoholic fermentation. Yeast. 2003;20(16): 1369–85.

39. Gadd GM, White C. Heavy metal and radionuclide accumulation and toxicity in fungi and yeasts. In: Poole RK, Gadd GM, editors. In Metal-Microbe Interactions. Oxford: IRL Press; 1989. pp. 19–38.

40. Gharieb MM, Gadd GM. Evidence for the involvement of vacuolar activity in metal(loid) tolerance: vacuolar-lacking and-defective mutants of *Saccharomyces cerevisiae* display higher sensitivity to chromate, tellurite and selenite. BioMetals. 1998;11(2): 101–6.

41. Cherest H, Davidian J, Benes V, Ansorge W, Surdin-kejan Y. Molecular Characterization of Two High Affinity Sulfate Transporters in Saccharomyces cerevisiae. Genet Soc Am. 1997;145(March): 627–35.

42. Tamás MJ, Labarre J, Toledano MB, Wysocki R. Mechanisms of toxic metal tolerance in yeast. In: Tamás MJ, Martinoia E, editors. Mol Biol Met Homeost Detoxif - From Mircobes to Man. Heidelberg, Springer Verlag; 2006. pp. 395–435.

43. Wysocki R, Tamás MJ. How *Saccharomyces cerevisiae* copes with toxic metals and metalloids. FEMS Microbiol Rev. 2010;34(6): 925–51.

44. Grant CM. Role of the glutathione/glutaredoxin and thioredoxin systems in yeast growth and response to stress conditions. Mol Microbiol. 2001;39(3): 533–41.

45. Li Y, Wei G, Chen J. Glutathione: a review on biotechnological production. Appl Microbiol Biotechnol. 2004;66(3): 233–42.

46. Zechmann B, Liou L-C, Koffler BE, Horvat L, Tomašić A, Fulgosi H, et al. Subcellular distribution of glutathione and its dynamic changes under oxidative stress in the yeast *Saccharomyces cerevisiae*. FEMS Yeast Res. 2011;11(8): 631–42.

47. Li Z-S, Szczypka M, Lu Y-P, Thiele DJ, Rea PA. The Yeast Cadmium Factor Protein (YCF1) Is a Vacuolar Glutathione S-Conjugate Pump. J Biol Chem. 1996;271(March 15): 6509–17.

48. Mendoza-Cózatl D, Loza-Tavera H, Hernández-Navarro A, Moreno-Sánchez R. Sulfur assimilation and glutathione metabolism under cadmium stress in yeast, protists and plants. FEMS Microbiol Rev. 2005;29(4): 653–71.

49. Paumi CM, Chuk M, Snider J, Stagljar I, Michaelis S. ABC transporters in *Saccharomyces cerevisiae* and their interactors: new technology advances the biology of the ABCC (MRP) subfamily. Microbiol Mol Biol Rev. 2009;73(4): 577–93.

50. Dhaoui M, Auchère F, Blaiseau P-L, Lesuisse E, Landoulsi A, Camadro J-M, et al. Gex1 is a yeast glutathione exchanger that interferes with pH and redox homeostasis. Mol Biol Cell. 2011;22(12): 2054–67.

51. Bidenne C, Blondin B, Dequin S, Vezinhet F. Analysis of the chromosomal DNA polymorphism of wine strains of *Saccharomyces cerevisiae*. Curr Genet. 1992;22(1): 1–7.

52. Mortimer RK. Evolution and variation of the yeast *(Saccharomyces)* genome. Genome Res. 2000;10(510): 403–9.

53. Borneman AR, Desany BA, Riches D, Affourtit JP, Forgan AH, Pretorius IS, et al. Whole-genome comparison reveals novel genetic elements that characterize the genome of industrial strains of *Saccharomyces cerevisiae*. PLoS Genet. 2011;7(2): e1001287.

54. Thomas D, Surdin-Kerjan Y. Structure of the HOM2 gene of *Saccharomyces cerevisiae* and regulation of its expression. Mol Gen Genet. 1989;217(1): 149–54.

55. Szczypka MS, Wemmie JA, Moye-Rowley WS, Thiele DJ. A yeast metal resistance protein similar to human cystic fibrosis transmembrane conductance regulator (CFTR) and multidrug resistance-associated protein. J Biol Chem. 1994;269(36): 22853–7.

56. Gueldry O, Lazard M, Delort F, Dauplais M, Grigoras I, Blanquet S, et al. Ycf1p-dependent Hg(II) detoxification in *Saccharomyces cerevisiae*. Eur J Biochem. 2003;270(11): 2486–96.

57. Prévéral S, Ansoborlo E, Mari S, Vavasseur A, Forestier C. Metal(loid)s and radionuclides cytotoxicity in *Saccharomyces cerevisiae*. Role of YCF1, glutathione and effect of buthionine sulfoximine. Biochimie. 2006;88(11): 1651–63.

58. Thorsen M, Perrone GG, Kristiansson E, Traini M, Ye T, Dawes IW, et al. Genetic basis of arsenite and cadmium tolerance in *Saccharomyces cerevisiae*. BMC Genomics. 2009;10: 105.

59. Eraso P, Martínez-Burgos M, Falcón-Pérez JM, Portillo F, Mazón MJ. Ycf1-dependent cadmium detoxification by yeast requires phosphorylation of residues Ser908 and Thr911. FEBS Lett. 2004;577(3): 322–6.

60. Swaney DL, Beltrao P, Starita L, Guo A, Rush J, Fields S, et al. Global analysis of phosphorylation and ubiquitylation cross-talk in protein degradation. Nat Methods. 2013;10(7): 676–82.

61. Fernandes L, Rodrigues-Pousada C, Struhl K. Yap, a novel family of eight bZIP proteins in *Saccharomyces cerevisiae* with distinct biological functions. Mol Cell Biol. 1997;17(12): 6982–93.

62. Sumner ER, Shanmuganathan A, Sideri TC, Willetts SA, Houghton JE, Avery S V. Oxidative protein damage causes chromium toxicity in yeast. Microbiology. 2005;151(Pt 6): 1939–48.

63. Ruotolo R, Marchini G, Ottonello S. Membrane transporters and protein traffic networks differentially affecting metal tolerance: a genomic phenotyping study in yeast. Genome Biol. 2008;9(4): R67.

64. Izquierdo A, Casas C, Muhlenhoff U, Lillig CH, Herrero E. *Saccharomyces cerevisiae* Grx6 and Grx7 Are Monothiol Glutaredoxins Associated with the Early Secretory Pathway. Eukaryot Cell. 2008;7(8): 1415–26.

65. Mesecke N, Spang A, Deponte M, Herrmann JM. A Novel Group of Glutaredoxins in the cis-Golgi Critical for Oxidative Stress Resistance. Mol Biol Cell. 2008;19: 2673–80.

66. Reinders A, Burckert N, Boller T, Wiemken A, De Virgilio C. *Saccharomyces cerevisiae* cAMP-dependent protein kinase controls entry into stationary phase through the Rim15p protein kinase. Genes Dev. 1998;12(18): 2943–55.

67. Fabrizio P, Pozza F, Pletcher SD, Gendron CM, Longo VD. Regulation of longevity and stress resistance by Sch9 in yeast. Science. 2001;292(5515): 288–90.

68. Cameroni E, Hulo N, Roosen J, Winderickx J, De Virgilio C. The Novel Yeast PAS Kinase Rim15 Orchestrates G0-Associated Antioxidant Defense Mechanisms. Genes Dev. 2004;3(4):

69. Myers LC, Gustafsson CM, Bushnell DA, Lui M, Erdjument-Bromage H, Tempst P, et al. The Med proteins of yeast and their function through the RNA polymerase II carboxy-terminal domain. Genes Dev. 1998;12(1): 45–54.

70. Sakurai H, Fukasawa T. Functional connections between mediator components and general transcription factors of *Saccharomyces cerevisiae*. J Biol Chem. 2000;275(47): 37251–6.

71. Harnpicharnchai P, Jakovljevic J, Horsey E, Miles T, Roman J, Rout M, et al. Composition and functional characterization of yeast 66S ribosome assembly intermediates. Mol Cell. 2001;8(3): 505–15.

72. Greenleaf AL, Kelly JL, Lehman IR. Yeast RPO41 gene product is required for transcription and maintenance of the mitochondrial genome. Proc Natl Acad Sci U S A. 1986;83(10): 3391–4.

73. Higgins VJ, Alic N, Thorpe GW, Breitenbach M, Larsson V, Dawes IW. Phenotypic analysis of gene deletant strains for sensitivity to oxidative stress. Yeast. 2002;19(3): 203–14.

74. Outten CE, Falk RL, Culotta VC. Cellular factors required for protection from hyperoxia toxicity in *Saccharomyces cerevisiae*. Biochem J. 2005;388(Pt 1): 93–101.

75. Bonawitz ND, Rodeheffer MS, Shadel GS. Defective Mitochondrial Gene Expression Results in Reactive Oxygen Species-Mediated Inhibition of Respiration and Reduction of Yeast Life Span. Mol Cell Biol. 2006;26(13): 4818–29.

76. Okada N, Ogawa J, Shima J. Comprehensive analysis of genes involved in the oxidative stress tolerance using yeast heterozygous deletion collection. FEMS Yeast Res. 2014;14(3): 425–34.

77. Zhu Y, Sun J, Zhu Y, Wang L, Qi B. Endogenic oxidative stress response contributes to glutathione over-accumulation in mutant *Saccharomyces cerevisiae* Y518. Appl Microbiol Biotechnol. 2015;99(17): 7069–78.

78. Hinnebusch AG. General and Pathway-specific Regulatory Mechanisms Controlling the Synthesis of Amino Acid Biosynthetic Enzymes in Saccharomyces cerevisiae. Vol. 21B, Cold Spring Harbor Monograph Archive. 1992. p. 319–414.

79. Düring-Olsen L, Regenberg B, Gjermansen C, Kielland-Brandt MC, Hansen J. Cysteine uptake by *Saccharomyces cerevisiae* is accomplished by multiple permeases. Curr Genet. 1999;35(6): 609–17.

80. Regenberg B, Düring-Olsen L, Kielland-Brandt MC, Holmberg S. Substrate specificity and gene expression of the amino-acid permeases in *Saccharomyces cerevisiae*. Curr Genet. 1999;36(6): 317–28.

81. Smith FW, Hawkesford MJ, Prosser IM, Clarkson DT. Isolation of a cDNA from *Saccharomyces cerevisiae* that encodes a high affinity sulphate transporter at the plasma membrane. Mol Gen Genet. 1995;247(6): 709–15.

82. Masselot M, De Robichon-Szulmajster H. Methionine biosynthesis in *Saccharomyces cerevisiae*. Mol Gen Genet. 1975;139(2): 121–32.

83. Thomas D, Surdin-Kerjan Y. Metabolism of sulfur amino acids in *Saccharomyces cerevisiae*. Microbiol Mol Biol Rev. 1997;61(4): 503–32.

84. Filetici P, Martegani MP, Valenzuela L, Gonzalez A, Ballario P. Sequence of the GLT1 gene from *Saccharomyces cerevisiae* reveals the domain structure of yeast glutamate synthase. Yeast. 1996;12(13): 1359–66.

85. Albers E, Laizé V, Blomberg A, Hohmann S, Gustafsson L. Ser3p (Yer081wp) and Ser33p (Yil074cp) are phosphoglycerate dehydrogenases in *Saccharomyces cerevisiae*. J Biol Chem. 2003;278(12): 10264–72.

86. Jaspers CJ, Gigot D, J. Penninckx M. Pathways of glutathione degradation in the yeast *Saccharomyces cerevisiae*. Phytochemistry. 1985;24(4): 703–7.

87. Adamis PDB, Panek AD, Eleutherio ECA. Vacuolar compartmentation of the cadmium-glutathione complex protects *Saccharomyces cerevisiae from mutagenesis*. Toxicol Lett. 2007;173(1): 1–7.

88. Ubiyvovk VM, Blazhenko O V, Gigot D, Penninckx M, Sibirny AA. Role of gamma-glutamyltranspeptidase in detoxification of xenobiotics in the yeasts Hansenula polymorpha and *Saccharomyces cerevisiae*. Cell Biol Int. 2006;30(8): 665–71.

